# *nanoFeatures:* a cross-platform application to characterize nanoparticles from super-resolution microscopy images

**DOI:** 10.1101/2024.02.12.579898

**Authors:** Cristina Izquierdo-Lozano, Niels van Noort, Stijn van Veen, Marrit M.E. Tholen, Francesca Grisoni, Lorenzo Albertazzi

## Abstract

Super-resolution microscopy and Single-Molecule Localization Microscopy (SMLM) are a powerful tool to characterize synthetic nanomaterials used for many applications such as drug delivery. In the last decade, imaging techniques like STORM, PALM, and PAINT have been used to study nanoparticle size, structure, and composition. While imaging has progressed significantly, often image analysis did not follow accordingly and many studies are limited to qualitative and semi-quantitative analysis. Therefore, it is imperative to have a robust and accurate method to analyze SMLM images of nanoparticles and extract quantitative features from them. Here we introduce *nanoFeatures*, a cross-platform Matlab-based app for the automatic and quantitative analysis of super-resolution images. *nanoFeatures* makes use of clustering algorithms to identify nanoparticles from the raw data (localization list) and extract quantitative information about size, shape, and molecular abundance at the single-particle and single-molecule levels. Moreover, it applies a series of quality controls, increasing data quality and avoiding artifacts. *nanoFeatures*, thanks to its intuitive interface is also accessible to non-experts and will facilitate analysis of super-resolution microscopy for materials scientists and nanotechnologies. This easy accessibility to expansive feature characterization at the single particle level will bring us one step closer to understanding the relationship between nanostructure features and their efficiency. https://github.com/n4nlab/nanoFeatures

## 1 Introduction

Nanoparticles have emerged as a promising tool in nanomedicine, owing to their remarkable physical and chemical properties [1, 2]. These physicochemical properties (*e.g*., size, shape, and presence and quantity of surface ligands) are key to selective delivery, via both passive [3], and active targeting [4]. These approaches hold the potential to mitigate side effects and reduce the required drug dosage [1].

Despite these advantages, there has been limited success in translating nanoparticles to clinical applications. Since 1995, when Doxil became the first FDA-approved liposome-based nano-drug for cancer treatment [5], only 31 formulations have been clinically approved, including the emergency approval for the COVID-19 vaccine [6]. The limited amount of approved nanoformulations shows that many open challenges still remain when translating the promising results of nanocarriers *in vitro* to a clinical application [7, 8]. Considering the vast number of nanoparticle formulations reported in literature, this evidence highlights the urgent need to promote the clinical approval of nanocarriers.

A current challenge is the lack of standardization in characterization methods, which results in a low degree of reliability and reproducibility in the nanomedicine literature [9]. Properly characterizing nanoparticles is a fundamental step in their potential application, as their physical and chemical properties can differ vastly at the nanoscale. Morphological features, like size or shape, heavily influence the nanoparticle properties, leading to unforeseen behavior or even toxicity in case of incorrect characterization [10, 2]. Currently available characterization techniques typically can only assess one property at a time, needing the integration of multiple techniques for a comprehensive characterization [2, 11]. Moreover, bulk measurement methods tend to average values and correspondingly mask the inherent heterogeneity of nanoparticles, which determines the effective distribution of nanoparticles to their site of action, and even their potential toxicity in biological media [10].

Recently, super-resolution microscopy [12] and, in particular, Single-Molecule Localization Microscopy (SMLM) [13] emerged as a powerful technique to improve nanoparticle characterization at the single-particle level. SMLM breaks the diffraction limit by computationally localizing individual fluorescent events separated in time and reconstructing super-resolved images based on these high-precision localizations. Therefore, SMLM images are created by stacking all the computed localization coordinates found in each frame of the acquisition movie, which allows researchers to visualize and analyze nanoparticles at the single-particle and single-molecule level with nanometer precision [13]. This technique has already shown its versatility in various research fields from cell biology [14, 15] to material characterization [16, 17], making it a powerful tool to address the challenges described above. For example, SMLM has been used to study and quantify protein corona formation on nanoparticles [18, 19], which revealed their heterogeneous nature. Furthermore, by combining SMLM with Transmission Electron Microscopy (TEM), multidimensional information to describe this heterogeneity could be obtained [20]. Moreover, SMLM has also allowed for the study of cell-nanoparticle interactions [21, 22]. Currently, research interest is shifting towards multiplexed images to correlate different modalities, which generate bigger datasets and complex data structures [23, 24, 25, 26, 27].

A crucial side of SMLM is represented by data analysis, as it is key to go from qualitative images to quantitative data. SMLM data analysis is based on: (1) processing the acquisition movie to obtain the localizations, through the fitting of the point-spread function (PSF) [28], (2) processing the localizations, for example, aligning or merging localizations [29], and finally (3) obtaining interpretable data [30] that we can use to measure size and morphology, thanks to the high spatial resolution of SMLM (20–50 nm) [13]. Modular platforms like Super-resolution Microscopy Analysis Platform (SMAP) allow the user to integrate many of the SMLM image analysis steps into a single software [31]. However, most of the available software resources are dedicated to biological imaging while materials and nanostructures lack tailored dedicated tools.

Here, we introduce *nanoFeatures*, an automated standalone MATLAB-based application dedicated to analyzing nanoparticles from SMLM datasets. The *nanoFeatures* app can process SMLM images, locate the nanoparticles, and automatically compute and display their features (Figure 1). After simply uploading the raw datasets and setting the right parameters, the user will receive a list of the key features for each individual nanoparticle, that can later be used in further analysis. By characterizing nanoparticles in a systematic and reproducible way, we aim to open the door for data-driven research, such as machine learning for property prediction or data mining. In what follows, we showcase an example application, whereby we analyse two different nanoparticle datasets: (1) dual color nanoparticles imaged with DNA-PAINT [32] and (2) triple color nanoparticles imaged with exchange PAINT [23].

**Figure 1:**
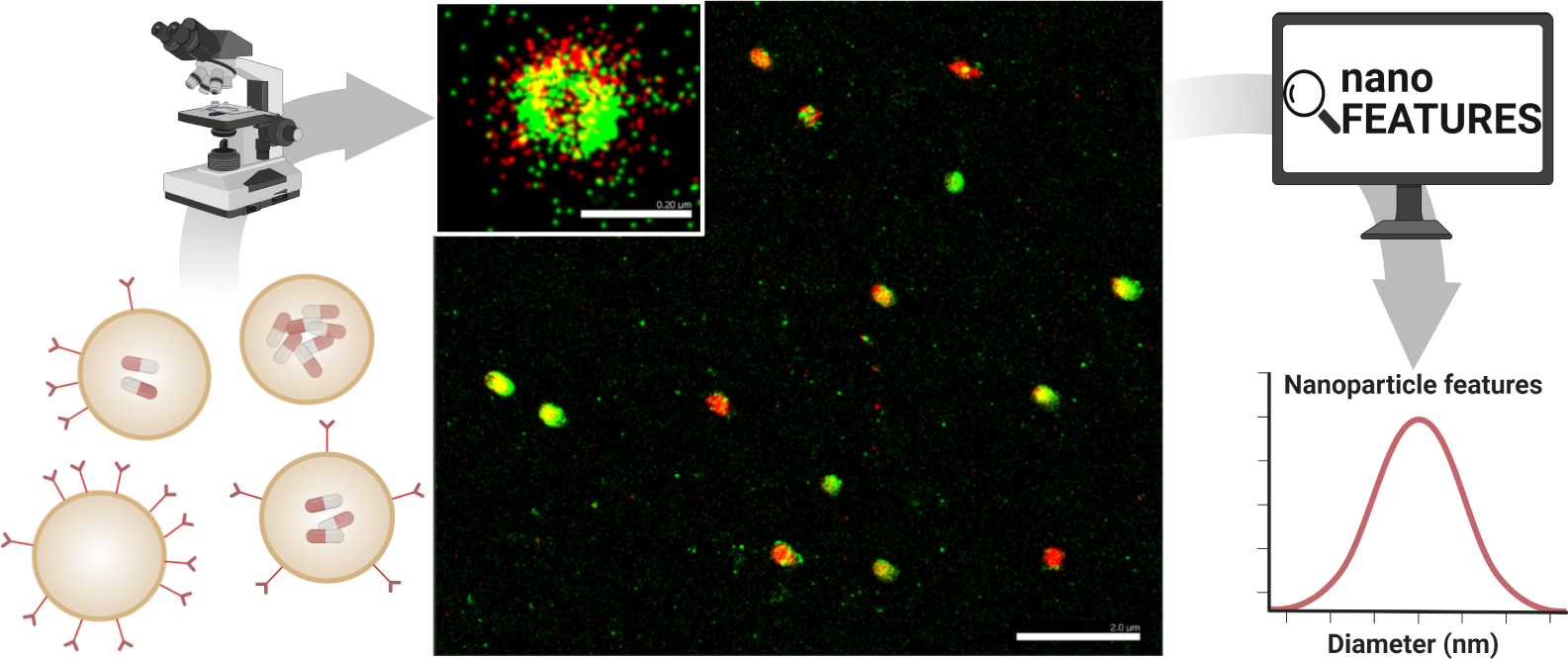
General nanoparticle analysis workflow. First, the nanoparticles are imaged in super-resolution microscopy. Then, the image data is sent to *nanoFeatures* to obtain nanoparticle features, such as size, shape, and number of binding sites. Scale bar 2 *μ*m.

## 2 Results and discussion

### 2.1 General procedure

A raw SMLM image is a list of coordinates for each of the fluorophore localizations found after fitting a Gaussian on the bright spots across the numerous frames in the acquisition movie (Supplementary Figure 1). An overview of the *nanoFeatures* algorithm is shown in Figure 2a.

**Figure 2:**
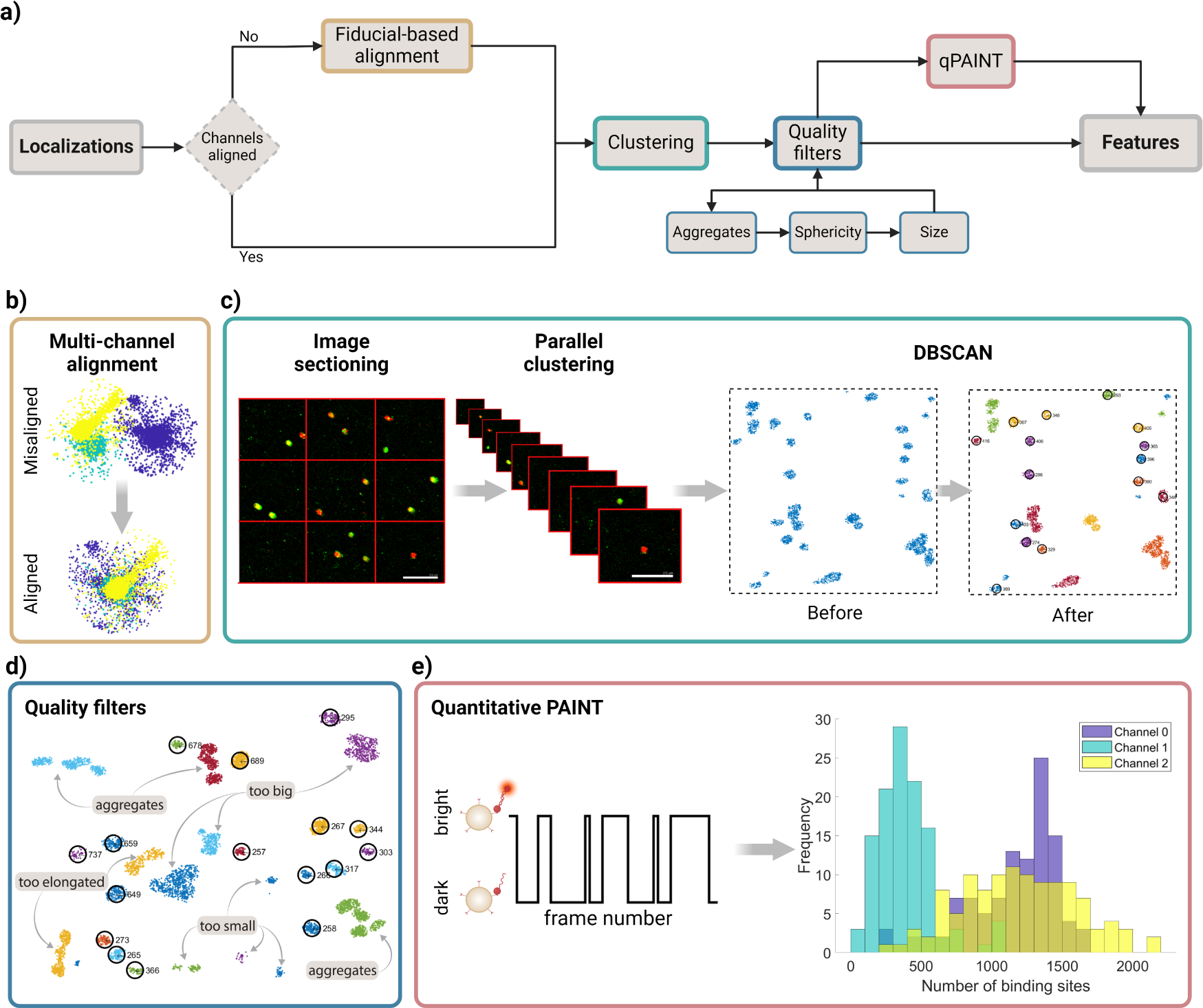
*nanoFeatures* algorithm. **a)** General workflow from the raw dataset to the calculation of the nanoparticle’s features. **b)** Example of the channel alignment results, in this case aligning fiducials from three different channels. **c)** Description of the image sectioning to send the different sections to parallel computing threads. MATLAB’s DBSCAN is used to identify clusters in the sections, an example of how this algorithm works is also shown. Scale bar 2 *μ*m. **d)** Quality filters applied to the identified clusters by DBSCAN. Aggregated, non-spherical, or off-size nanoparticles are filtered in this step. Selected clusters are circled in black and given an ID number. **e)** qPAINT scheme, the frames in which the single fluorophores are on or off are counted, obtaining the dark and bright times, and are used to infer the total number of binding sites.

The *nanoFeatures* general workflow is the following (Figure 2a):

1. User input: several inputs are required from the user, which will determine the following steps of the analysis. The steps are as follows:
  a. *File input*. The user inputs the file, as specified in the User Input subsection, containing the list of localization coordinates (Supplementary Figure 1). Multiple color channels can be input as a single file or as multiple files, corresponding to either simultaneous multi-color imaging (e.g. 2or 3-color STORM) or sequential multiplexing (e.g. exchange-PAINT). To increase the throughput, users can select multiple files to be processed under the same settings in batch analysis.
  b. *Input type selection*. After inputting the file, the user must select the input type (ONI, NIKON, or ThunderSTORM), based on the microscope or software used to obtain the images.
  c. *Workflow and parameter specification*. Within the Graphical User Interface (GUI), users can specify the desired workflow for data processing. For example, when analyzing samples acquired sequentially, the user should activate the channel alignment option. Furthermore, users can define the parameters for filtering and feature extraction, and have the option to include the quantitative PAINT (qPAINT) analysis. [33].
2. Read and pre-process file(s): Once the input, workflow, and parameters have been defined, the data is pre-processed into a single list of localizations, containing all color channels.
3. Channel alignment *(optional)* (Figure 2b): If specified, the drift between different color channels is corrected before clustering.
4. Clustering (Figure 2c). The processed localizations are then used as the input of a clustering algorithm, which will identify potential nanoparticles, using the vicinity information.
5. Quality filters (Figure 2d): A series of quality filters are applied to exclude clusters of localizations not identified as single nanoparticles.
6. qPAINT *(optional)* (Figure 2e): Quantifies the target ligands count based on the spatiotemporal response within each cluster.
7. Generate output: Finally, *nanoFeatures* will generate an output CSV file containing the features corresponding to each detected nanoparticle in the original SMLM image. This file, together with the plots prompted during the execution, will be saved in a “results” folder in the user’s Matlab path.

In addition, SMLM images often have high background noise, leading to prolonged computing time for clustering analysis and complicating the automated identification of nanoparticles. For this reason, we advise pre-processing the image before using *nanoFeatures*, for example, by applying a density filter, as seen in Supplementary Figure 2. In this specific case study, we used the thunderSTORM plug-in for ImageJ [34], with the settings outlined in the Density Filter subsection in Materials and Methods. Moreover, a description of the datasets used for this case study can be found in the Datasets subsection in Materials and Methods.

The following sections provide a detailed description of each of these steps.

### 2.2 User input

*nanoFeatures* accepts three distinct data structures, corresponding to outputs from various commercial microscopes or open-source image pre-processing software. The app extracts the pixel localizations (X, Y), the frame of detection, and the associated color channel. The three supported input file types are:

1. **NIKON** [35]: TXT file format. The app reads the file as a table, identifying headers named “Channel Name”, “X”, “Y” and “Frame”.
2. **ONI** [36]: Comma-separated values (CSV) file format. The app reads the file as a matrix, considering the 1^st^ column as the channel name, 2^nd^ column as frame number, and the 3^rd^ and 4^th^ columns as X and Y coordinates, respectively.
3. **ThunderSTORM** [34]: CSV file format. The app reads the file as a table, identifying headers named “Channel”, “Frame”, “x [nm]” and “y [nm]”.

Extension to further formats is envisioned in the future to follow the evolution of the different formats used in the community.

As an example, we showcase the workflow and parameter specification for the analysis of the file “210714 COOH 200 loc2 merge.csv”, which can be found in our online repository under the files “Datasets/dualColor/COOH 200nm 100dist 50neighbors”. The workflow and parameter selection can be seen in Figure 3a-c and the results preview given by *nanoFeatures* within the GUI is seen in Figure 3d.

**Figure 3:**
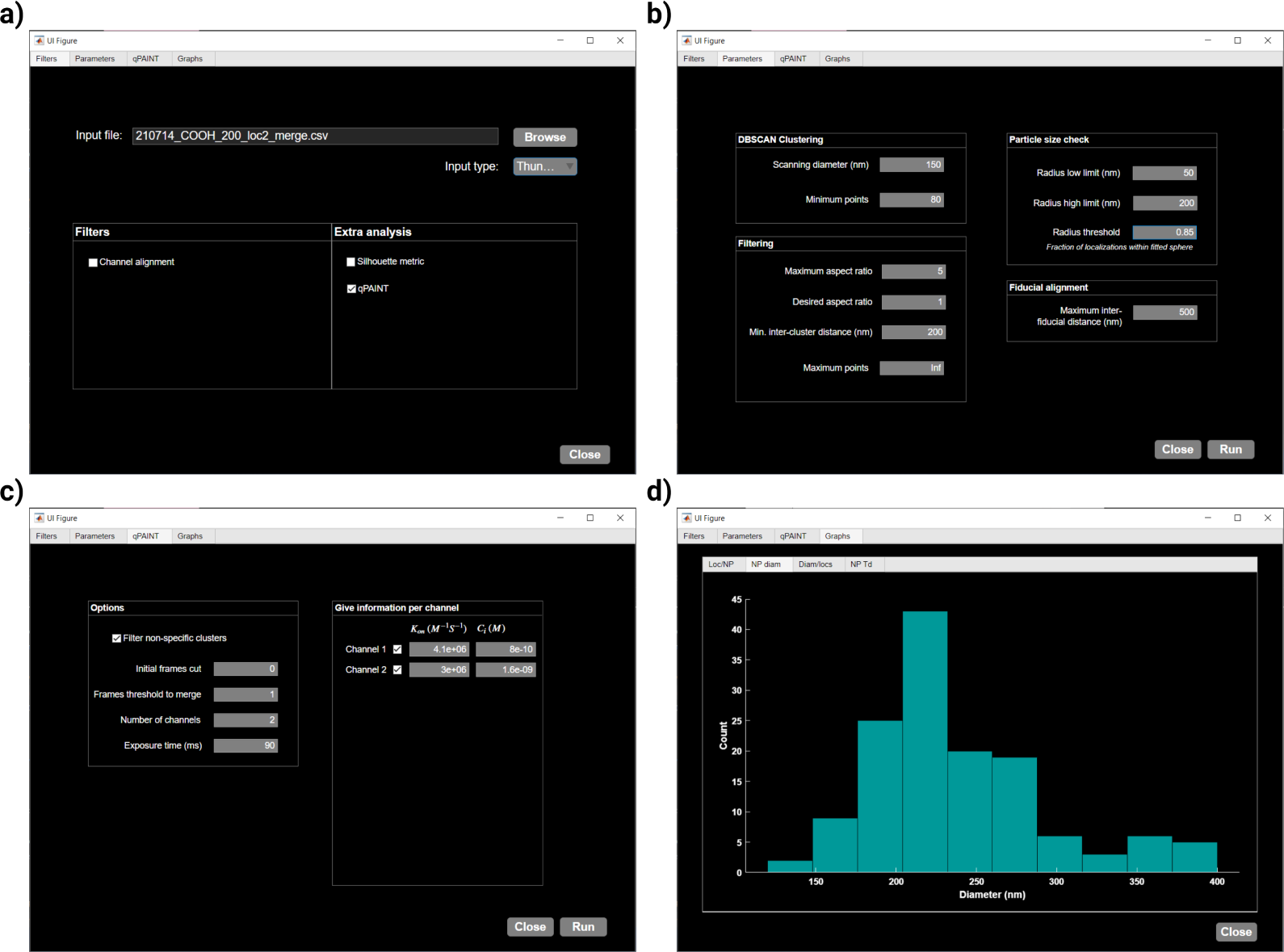
Case study using the *nanoFeatures* Graphical User Interface (GUI). **a)** Filters tab to input the file(s) and select the desired filters, **b)** parameters tab to input the specific parameters used to analyze the files, **c)** qPAINT tab to input the parameters for the qPAINT analysis, if the checkbox is selected, and **d)** graphs tab to show a preview of the results.

### 2.3 Channel alignment

SMLM images are often acquired sequentially, one color at a time, such as through exchange-PAINT [23]. The resulting data is stored in multiple localization files, one for each color. Since the image acquisition is done at different time points, there is a drift between the coordinates of different channels that we need to correct, as shown in Figure 2b. To achieve this, we use fiducial markers —objects consistently visible in all color channels that serve as reference points. In this study, we use gold nanorods as fiducial markers.

For this reason, upon clicking the “Run” button and only if the “channel alignment” checkbox is selected, *nanoFeatures* prompts the user to input files containing the fiducial localizations for each respective color channel.

To correct the temporal drift, first, the fiducial localizations are clustered to obtain their centroid. Then, *nanoFeatures* finds the *n-channels* nearest neighbors within a limited distance, to avoid fiducial mismatching. Finally, the average drift distances for each fiducial match between sequential files are calculated, after which the coordinates are corrected and the files are merged.

### 2.4 Clustering

After aligning all localizations from the different color channels and concatenating the files into a single matrix, the localizations are ready for clustering. This is a crucial step needed to identify individual nanoparticles and assign all the detected molecules to a specific nanoparticle or discard them as background.

To speed up computational time, the image is divided into nine sections according to their coordinates. Then these sections are processed simultaneously in *nCores −* 1 *<*= 9 parallel computing threads, running the DBSCAN algorithm [37], as described in Figure 2c. This parallelization step significantly reduces the execution time, while preventing the app from potential collapse due to the generation of massive data structures required to link all localizations within a background-dense image (Supplementary Figure 3). Note that at least one core remains available for other computer processes.

DBSCAN requires two parameters: (1) the radius in which adjacent localizations are considered as neighbors, and (2) the minimum number of points in this neighborhood required to form a cluster. These are introduced by the user, and they require optimization for each experiment and sample. To do this, the user can select the Silhouette checkbox in *nanoFeatures* to plot the Silhouette coefficient [38] for each cluster, as in Supplementary Figure 4f. However, this analysis is computationally expensive, hence it is not recommended to use with large files or during batch analysis.

Then, *nanoFeatures* plots the reconstructed complete image (all nine sections), with the identified clusters by DBSCAN, which are depicted by different colors as shown in Figure 4a.

**Figure 4:**
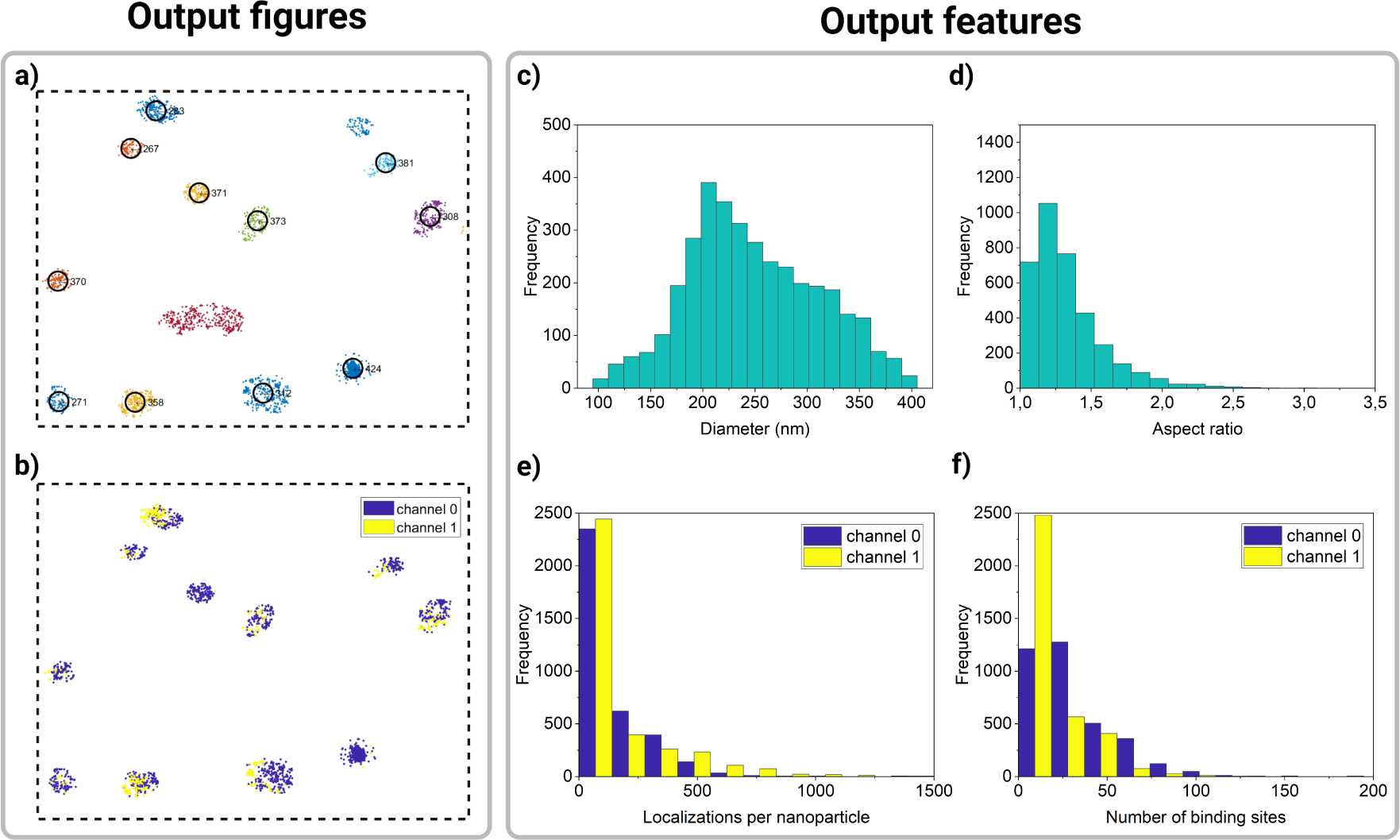
Selection of different figures obtained from *nanoFeatures*. **a)** DBSCAN identified clusters and nanoparticles selected by the quality filters. **b)** Selected nanoparticles colored based on the channel that each of the localizations was identified in. **c-f)** Histograms showing the distribution for a few of the different features obtained from *nanoFeatures*. Graphs generated in MATLAB (a-b) and OriginLab (c-f), based on a 200 nm spherical nanoparticles sample.

### 2.5 Quality filters

Although DBSCAN can identify clusters in images, and discard some localizations based on userdefined parameters, some quality filters are necessary to discard unwanted signals. For example, large aggregates or debris need to be discarded. Moreover, a strong background signal may be detected by DBSCAN as a particle. Therefore, to ensure the identified clusters are indeed nanoparticles, the data undergoes the following quality filters: (1) The sphericity filter removes clusters that are too elongated, or not elongated enough, depending on the sample. (2) The size filter removes clusters that are too big or too small. And (3) the aggregates filter removes clusters that are too close to each other (Figure 2d).

To do this, each filter: (1) fits an ellipse on the clusters to obtain the aspect ratio. Then, (2) sets a random starting radius and adjusts it until a user-defined percentage of the cluster localizations fit in it. Finally, (3) ensures the distance between each cluster centroid is at least the user-defined distance.

As a result of the quality filters, some clusters identified by DBSCAN are discarded, and only potential nanoparticles remain, marked by a black circle and an ID number in Figure 4a. Additionally, *nanoFeatures* plots the selected nanoparticles from all nine sections, color-coded based on the localization’s channel. This way, users can identify co-localizing ligand populations on a nanoparticle (Figure 4b).

At this point, within the GUI “Graphs” tab, *nanoFeatures* generates histograms providing an overview of the nanoparticles’ characterization (Figure 3d). For a more comprehensive analysis, users can plot the features from the generated CSV file, as showcased in Figures 4c-f. For instance, these plots show features from a sample of 200 nm spherical nanoparticles.

Figure 4c shows a wide distribution of nanoparticle diameters, peaking around 200 nm. Similarly, Figure 4d shows the aspect ratio for the same sample, with its distribution mostly encompassed between 1 and 1.5, where 1 is a perfect sphere. These results align with the heterogeneous nature of nanoparticles, as described in literature [10].

### 2.6 qPAINT

Finally, if the sample was obtained following a DNA-PAINT protocol, users have the option to perform a quantitative PAINT (qPAINT) analysis [33] (Figure 2e). First, the qPAINT filter generates a binary time trace for each identified cluster: bright (1), for each frame in which the fluorophores are active, or dark (0) when they are inactive.

Next, the user introduces the minimum number of frames that a cluster needs to be on (not dark), for the localizations to be merged into a single event. This number, the “frames threshold to merge”, needs to be optimized per sample, and it prevents false dark times [39]. The duration of each dark time is then calculated by taking the derivative of the nanoparticle time traces and finding the difference between consecutive negative and positive changes. Clusters formed by non-specific localizations can be filtered by checking the corresponding checkbox within the GUI. Clusters will then be removed if their bright times do not comprise at least 50% of the total imaging time between their first and last binding event.

Further, *nanoFeatures* constructs a cumulative distribution function (CDF) of the dark times for each remaining cluster, which is fitted with eq. 1, to obtain its mean dark time (*τ*_*d\**_), where P represents the probability of a binding event at time t after a previous binding event (Supplementary Figure 5). Moreover, clusters are filtered on their CDF shape. When 90% of dark times are smaller than 10% of the longest dark time, the cluster is discarded.

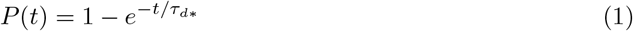

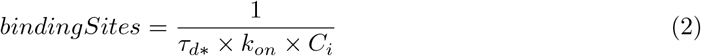

Then, *τ*_*d\**_, in milliseconds, is used in eq. 2 to determine the number of binding sites per cluster (Figure 4f). The association constant (*k*_*on*_) for each docking-imager pair and the imager concentration (*C*_*i*_) are experiment-dependent.

Lastly, *nanoFeatures* will create a folder named “results” on the user’s current Matlab path. This folder contains, for each file analyzed, most of the figures plotted by Matlab (Supplementary Figure 4) and a CSV file containing the number of binding sites, statistics of the bright and dark times, and R-squared for the exponential fit. A detailed list of all the features exported by *nanoFeatures* can be found in Supplementary Table 2.

## 3 Conclusion

*nanoFeatures* aims to fill the gap in standardizing nanoparticle characterization, particularly by analyzing super-resolution microscopy images. *nanoFeatures* achieves this by integrating multiple SMLM techniques, commercial microscopes and hardware set-ups, and diverse file formats. These can all be analyzed within the same algorithm, generating features organized in a unified data structure.

Moreover, despite being a Matlab-based application, the standalone version can be installed via MATLAB Runtime (without the need for a license), on multiple platforms like Linux, macOS, and Windows.

This way, the many files generated from diverse experiments, can be directly used in further analyses, minimizing or eliminating the need for extensive data pre-processing. For instance, employing machine learning to analyze various nanoparticle samples could provide valuable insights into the interrelationships of these features, thus aiding in the future design of nanoparticle formulations.

## 4 Materials and methods

### 4.1 Datasets

The results were obtained by analyzing two different datasets. The main dataset used depicts two different antibodies on nanoparticles surface, imaged simultaneously with two different fluorophores (two color channels). The structure of this dataset consists of a list of localizations with information on their specific coordinates, frame number, and photon intensity. This data was published by Marrit M.E. Tholen et al. (2023) [17]. To test the channel alignment filter, a DNA PAINT dataset consisting of both 300 nm polystyrene particles with a conjugated ssDNA and 40nm gold nanorod fiducials was used. The sample was measured at three different time points in multiple fields of view. These datasets, together with the *nanoFeatures* results, can be found in our online repository in Zenodo.

Figures 2b and 2e are based on the exchange PAINT files in the “tripleColor” folder. Figures 4a-b are based on the file *“210714 COOH 200 loc2 merge.csv”* from Marrit Tholen’s dataset and Figures 4c-f are based on the combination of all files from sample *“COOH 200nm”*. These files can be found in the “dualColor” folder.

### 4.2 Drift correction

Fiducial markers can be used for drift correction due to the fact that they are stationary and permanently in an ‘on’ state. This was done in thunderSTORM, using a ‘Minimum marker visibility ratio’ of 0.5, which means they should be on for at least half the total frames to be considered as a fiducial marker. The ‘Max distance [units of x,y]’, or the lateral tolerance for identification as a marker is usually set to 60.0 nm. The ‘Trajectory smoothing factor’ is set to 0.5. When no fiducial markers are used, a cross-correlation function can be used.

### 4.3 Fiducial localization files

Similarly, fiducials can also be identified by making use of the fact that they are permanently in an ‘on’ state. By applying an extreme density filter of 9000 minimum neighbors in a 50 nm radius, all nanoparticle clusters are filtered out and only the fiducials remain. This way, we can obtain the fiducial coordinates to be used within the channel alignment option.

### 4.4 Density filter

Prior to analyzing the images in *nanoFeatures*, we used the thunderSTORM plug-in for ImageJ [34] to remove the background noise. The settings are 100 nm radius and 60 minimum neighbors for sample COOH 100, 100 nm radius and 50 minimum neighbors for sample COOH 200, 100 nm radius and 100 minimum neighbors for sample COOH 300, and 100 nm radius and 50 minimum neighbors for sample NH2 200. Table 1 shows the files that were filtered with different settings.

**Table 1:**
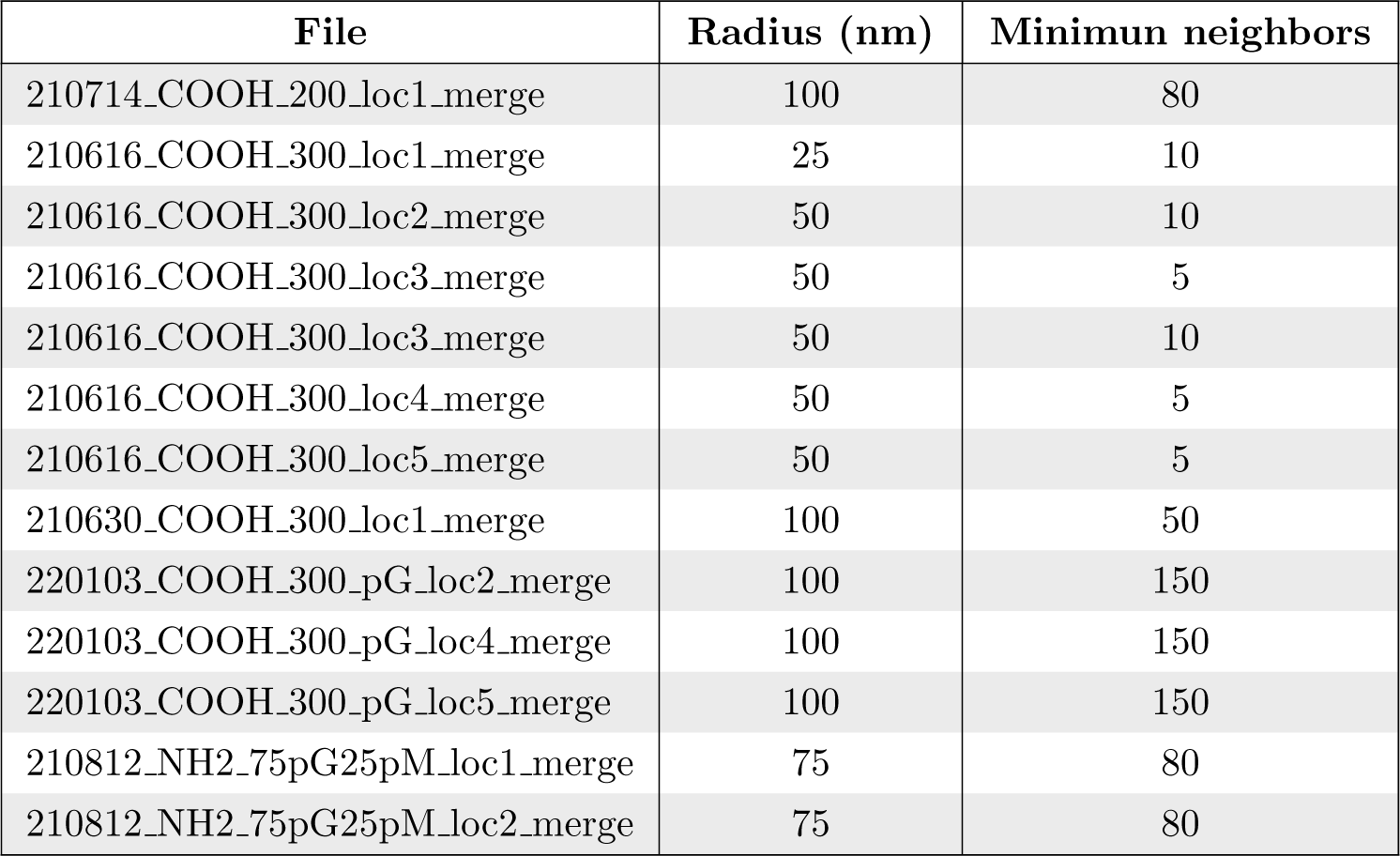
Specific density filter settings.

### 4.5 Hardware and software

In this work, *nanoFeatures* was run on MATLAB v. 2022b [40], on an AMD Ryzen 9 5900X 12Core Processor 3.70 GHz with 32GB RAM and a 64-bit Windows 10 Enterprise machine (Version 10.0.19045 Build 19045).

## 5 Acknowledgements

The authors would like to thank Pietro del Canale and Roger Riera Brillas for the original scripts that *nanoFeatures* is based on, and for engaging in algorithm discussions.

## 6 Author Contributions

*Conceptualization*: CIL, LA, and FG. *Data curation:* MMET, SvV, NvN, and CIL. *Formal analysis:* CIL, MMET, SvV, and NvN. *Methodology:* CIL, FG, and LA. *Software:* CIL, and NvN. *Visualization:* CIL. *Writing - original draft:* CIL. *Writing - review and editing:* all authors. All authors have approved the final version of the manuscript.

## 7 Code availability

The code can be found on GitHub https://github.com/n4nlab/nanoFeatures, and the datasets used in Zenodo https://zenodo.org/records/10610875

## 8 Declaration of interest

Biorender was used to design the figures in this work. The authors used ChatGPT as a tool to improve readability and clarity of the manuscript the authors reviewed and edited the content as needed and take full responsibility for the content of the publication.

## Notes

https://github.com/n4nlab/nanoFeatures

https://zenodo.org/records/10610875

### Competing Interest Statement

The authors have declared no competing interest.

### Summary of Updates

The image was resolution improved in this version.

