## Supplementary Information for "*nanoFeatures:* a cross-platform application to characterize nanoparticles from super-resolution microscopy images"

### Supplementary information
### *nanoFeatures*

Cristina Izquierdo-Lozano, Niels van Noort, Marrit M.E. Tholen, Stijn Van Veen, Francesca Grisoni and Lorenzo Albertazzi

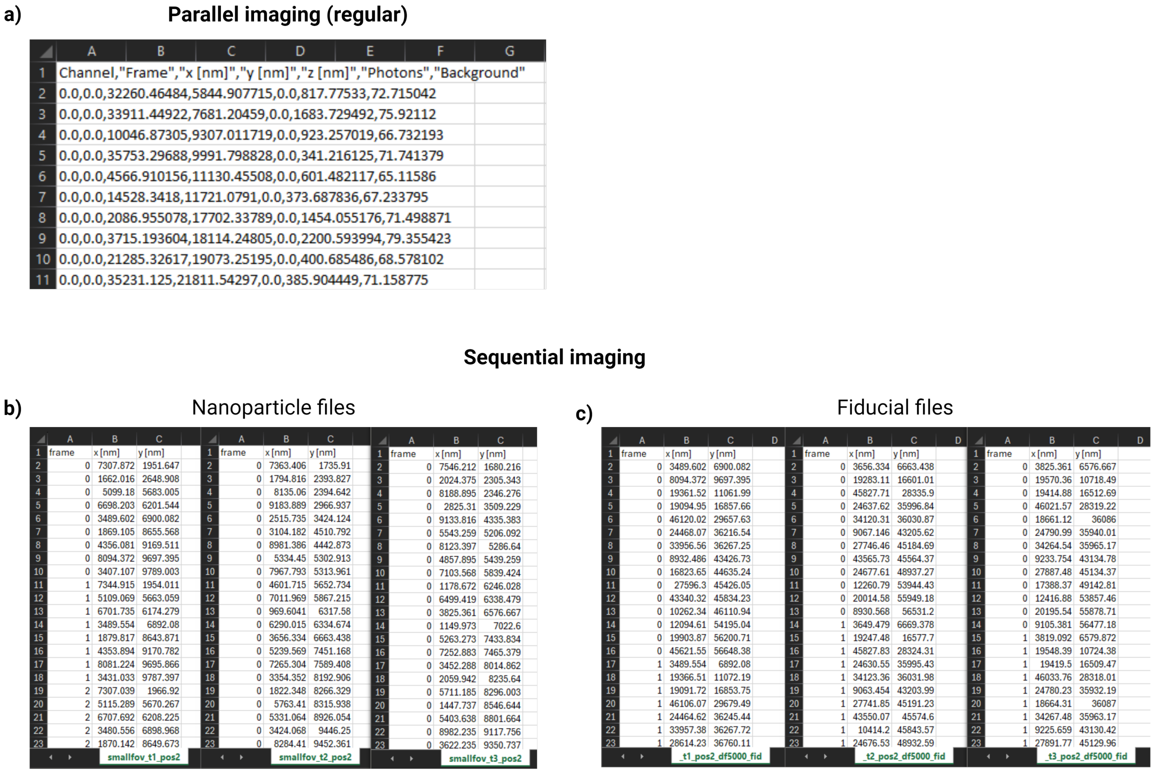


*Figure 1:* Raw data example of nanoparticles imaged in super-resolution microscopy (DNA-PAINT), **a)** all color channels imaged in the same acquisition and **b)** three different color channels imaged sequentially, which has to include **c)** the corresponding files containing the fiducial localizations.

*Table 1:* Description of the input parameters required by *nanoFeatures*.

| **Parameter** | **Description** |
|---|---|
| ***Filters tab*** | |
| Input file | Files(s) to be analyzed by *nanoFeatures*. |
| Input type | Microscope or software the files were obtained from. Current options are Nikon (N-STORM), Oxford Nanoimager (ONI) and ThunderSTORM (ImageJ plugin). |
| Channel alignment | Checkbox to align the different channel colors in the case of exchange PAINT (sequentially imaging each color instead of simultaneously). This filter is based on fiducials, then the user would need to first input the different files (one per color) and then nanoFeatures will ask to input the files containing the fiducial localizations. This filter doesn't admit batch analysis. |

| | |
|---|---|
| Silhouette metric | Checkbox to calculate the Silhouette coefficient for each cluster found by DBSCAN. This metric can be used while optimizing the parameters and measures how well the nanoparticles are clustered. The closer to 1 the better. The calculation of this metric is computationally expensive if the images are too dense. |
| qPAINT | Checkbox to perform the qPAINT analysis. That is, to calculate the actual number of molecular targets based on the number of localizations and binding kinetics. |
| ***Parameters tab*** | |
| *DBSCAN clustering* | |
| Scanning diameter | Diameter, in nanometers, used by DBSCAN to follow the density of points. Generally, it is the same as the size of the nanoparticles in the image. However, if these are too dense, it is better to set a smaller diameter. |
| Minimum points | Minimum number of localizations that have to be contained in the scanning diameter to be considered a full cluster or part of a bigger cluster. Relates to the nanoparticle's density. |
| *Filtering* | |
| Maximum aspect ratio | Maximum ellipticity of the nanoparticles, being 1 a perfect sphere. |
| Desired aspect ratio | Theoretical shape of the nanoparticles being analyzed. |
| Min. inter-cluster distance | Minimum separation, in nanometers, between nanoparticles. Generally, at least the same distance as their diameter. |
| Maximum points | Maximum number of points for a cluster to be considered a nanoparticle. |
| *Particle size check* | |
| Radius low limit | Lowest radius size, in nanometers, for a cluster to be considered a nanoparticle. |
| Radius high limit | Highest radius size, in nanometers, for a cluster to be considered a nanoparticle.. |
| Radius threshold | Fraction of the localizations within the radius. Sometimes some isolated localizations in a cluster might be further away from the actual radius and would be better to exclude them. |
| *Fiducial alignment* | |
| Maximum inter-fiducial distance | Maximum separation, in nanometers, for fiducials in different channels to be considered the same. Avoids mismatching. |
| ***qPAINT tab*** | |
| *Options* | |
| Filter non-specific clusters | Checkbox to filter out clusters that are not present for at least 50% of the imaging time. |
| Initial frames cut | Number of frames to cut from the beginning of the imaging movie. |
| Frames threshold to merge | Allowed gap in between binding events for them to be still considered consecutive. |
| Number of channels | Number of laser channels (colors) present in the analyzed images. |
| Exposure time | Acquisition's exposure time, in milliseconds. |
| *Give information per channel* | |
| $(k_{on})$ | Association constant for each docking-imager pair, determined experimentally. |
| $(C_i)$ | Imager concentration. |
| ***Graphs tab*** | |
| Loc/NP | Number of localizations per nanoparticle histogram. |
| NP diam | Nanoparticle's diameter (size) histogram. |
| Diam/locs | Nanoparticle's localizations compared to their size scatterplot. |
| NP Td | True dark time per nanoparticle histogram. |

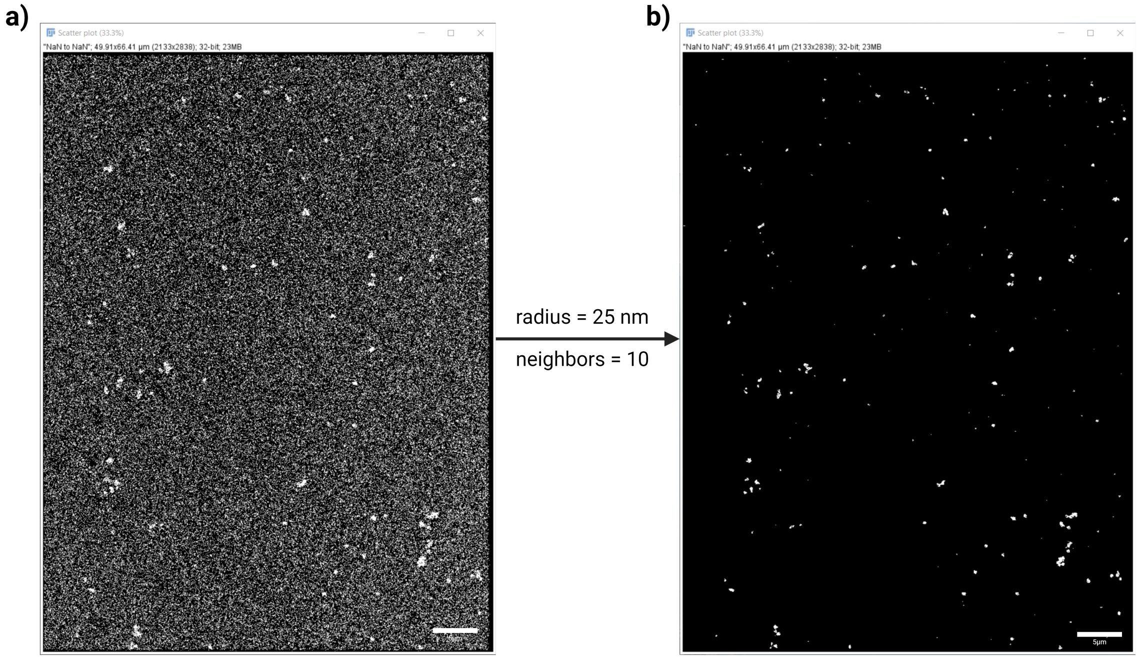


*Figure 2:* Before and after example of a density-filtered super-resolution microscopy image. In this case, an image of 300 nm nanoparticles was filtered by density with a radius of 25 nm and 10 minimum number of neighbors. Scale bar: 5µm.

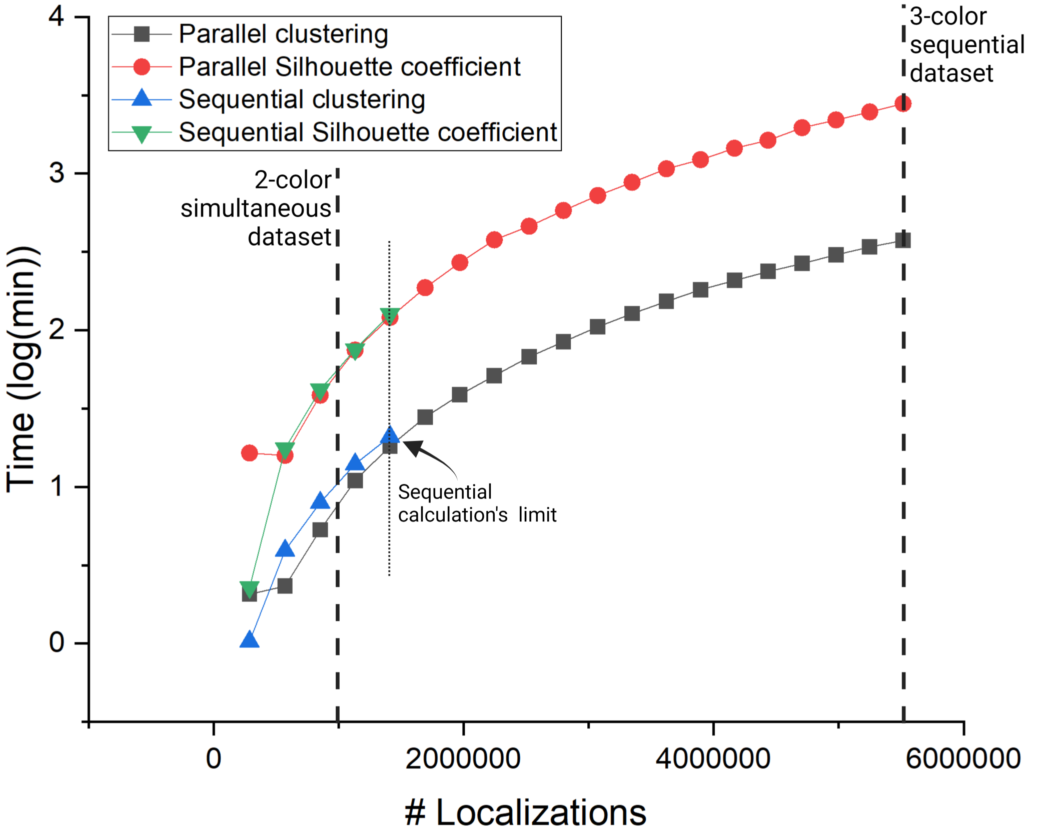


*Figure 3:* Time analysis of the *nanoFeatures* execution, comparing between parallel and sequential execution of the nine sections and with or without the Silhouette analysis. Note that sequential analysis is only apparent for the lower number of localizations due to rapidly increasing computational costs and memory requirements, resulting in the execution breaking.

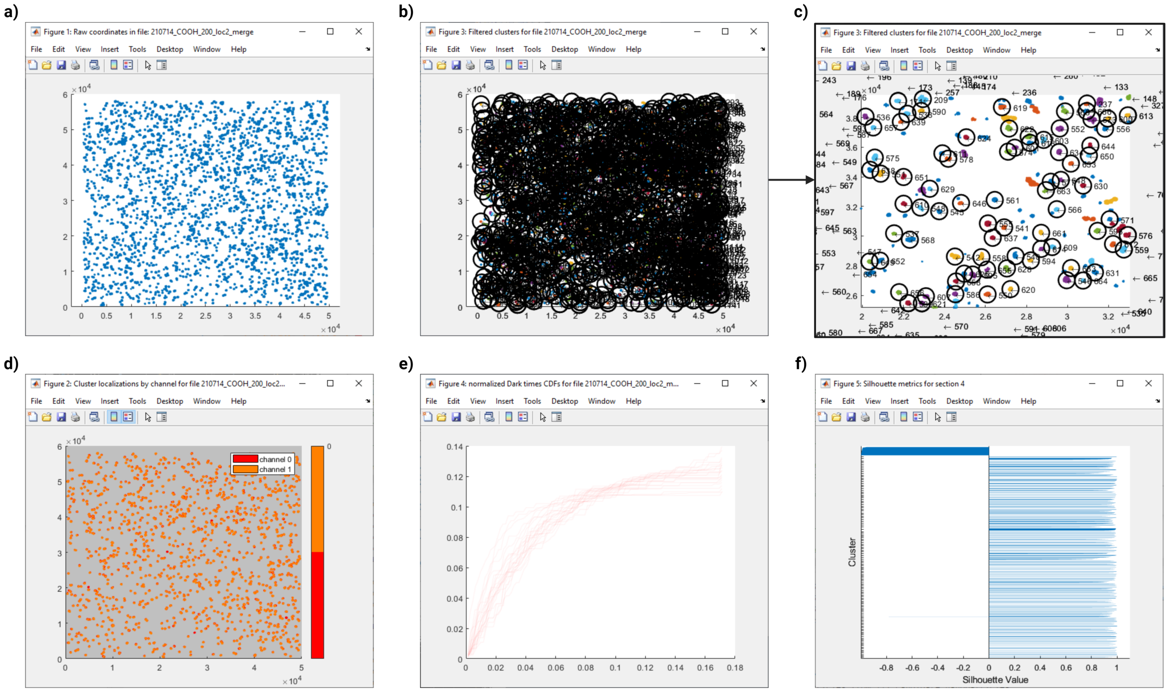


*Figure 4:* Example of *nanoFeatures* output figures. In this case, for DNA-PAINT dual-color 200nm nanoparticles. **a**) Raw coordinates, plotted directly from the localization list input by the user. **b)** Identified nanoparticles by DBSCAN, after going through the quality filters and **c)** the zoom in. Colored clusters are the ones identified by DBSCAN and the circled clusters are selected by the quality filters. **d)** Selected nanoparticles colored based on the channel each localization was found in. **e)** normalized Cumulative Distribution Function (CDF) of each nanoparticle's dark times. **f)** Silhouette metric for each of the identified clusters by DBSCAN, from -1 (not clustered) to 1 (perfectly separated cluster). Note that the -1 group corresponds to the background localizations, discarded by DBSCAN. Group ID (given by DBSCAN) can be found by clicking in any of the localizations colored in *nanoFeature*'s figure 3 (**b**) in this figure).

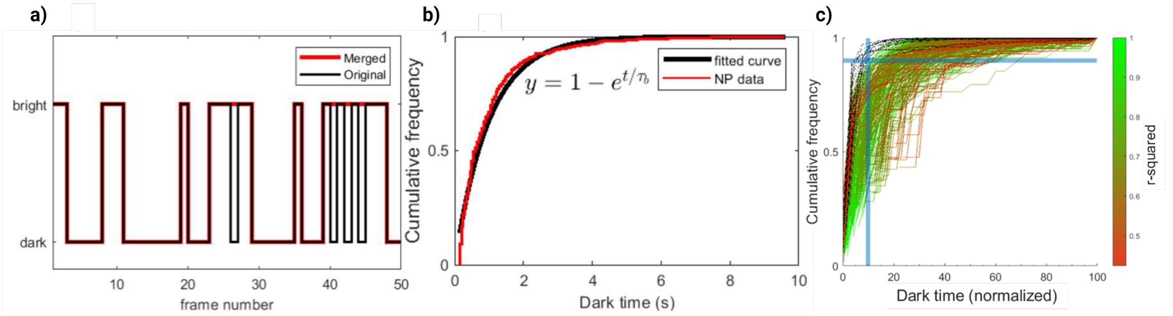


*Figure 5:* qPAINT filter **a)** The first 50 frames of a binary particle time trace (black). Consecutive binding events (when the particle is 'bright') with a gap of up to three frames are merged into a single event, resulting in the merged time trace shown in red. **b)** Example of the dark time CDF for a cluster (red) superimposed with its fitted curve (black), computed using equation 1. **c)** All normalized CDFs for a measurement, colored by their R-squared value from a bad fit (red) to a good fit (green). CDFs with an unexpected shape (black) that go above the threshold (blue cross) are filtered out.

*Table 2:* Description of the features obtained by *nanoFeatures*.

| **Feature** | **Description** |
|---|---|
| Diameter | The diameter of the cluster as determined by ellipse fit-ting from *nanoFeatures*. |
| Aspect ratio (shape) | The aspect ratio (starting at one) of the ellipse fitted over the cluster by *nanoFeatures*. |
| x-coordinate | The x-coordinate of the cluster center determined by *nanoFeatures*. This can be used for the reconstruction of cluster centers. |
| y-coordinate | The y-coordinate of the cluster center determined by nanoFeatures. This can be used for the reconstruction of cluster centers. |
| Cluster localizations | The total number of localizations in the cluster. |
| ***For each channel*** | |
| Channel localizations | The number of localizations of the cluster that are in the corresponding channel. |
| True mean dark time | The mean dark time of the localizations in the corresponding channel as determined by qPAINT analysis through CDF fitting. |
| R-squared | The goodness of fit of the CDF fitting expressed in R-squared. |
| Mean dark time | The mean value of the dark times in the corresponding channel calculated during qPAINT calculations. |
| Median dark time | The median value of the dark times in the corresponding channel calculated during qPAINT calculations. |
| SD dark time | The standard deviation of the dark times in the corresponding channel calculated during qPAINT calculations. |
| Mean bright time | The mean value of the bright times in the corresponding channel calculated during qPAINT calculations. |
| Median bright time | The median value of the bright times in the corresponding channel calculated during qPAINT calculations. |
| SD bright time | The standard deviation of the bright times in the corresponding channel calculated during qPAINT calculations. |
| Target count | The number of binding sites for the corresponding channel as determined by qPAINT calculations using the true mean dark time and user-set parameters for acquisition frame rate and association constant ($k_{on}$). |
